# Numerical Simulation of Concussive-generated Cortical Spreading Depolarization to Optimize DC-EEG Electrode Spacing for Non-invasive Visual Detection

**DOI:** 10.1101/2021.04.08.438969

**Authors:** Samuel J. Hund, Benjamin R. Brown, Coline L. Lemale, Prahlad G. Menon, Kirk A. Easley, Jens P. Dreier, Stephen C. Jones

**Affiliations:** CerebroScope, the dba entity of SciencePlusPlease LLC, 1936 Fifth Avenue, Pittsburgh, PA 15219-5544, USA; SimulationSolutions, LLC, 5707 Jackson St., Pittsburgh, PA, 15206, USA; Center for Stroke Research, Charité – Universitätsmedizin Berlin, corporate member of Freie Universität Berlin, Humboldt-Universität zu Berlin, and Berlin Institute of Health, Berlin, Germany; Department of Experimental Neurology, Charité – Universitätsmedizin Berlin, corporate member of Freie Universität Berlin, Humboldt-Universität zu Berlin, and Berlin Institute of Health, Berlin, Germany; Electrical & Computer Engineering, SYSU-CMU Joint Institute of Engineering, Sun Yat-sen University - Carnegie Mellon University (SYSU-CMU), 5000 Forbes Ave., Pittsburgh, PA, 15213, USA, Department of Bioengineering, University of Pittsburgh, 302 Benedum Hall, Ohara St., Pittsburgh, PA, 15213, USA; Department of Biostatistics and Bioinformatics, Rollins School of Public Health, Emory University, 1518 Clifton Road NE, Atlanta, GA, 30322, USA; Department of Neurology, Charité – Universitätsmedizin Berlin, corporate member of Freie Universität Berlin, Humboldt-Universität zu Berlin, and Berlin Institute of Health, Berlin, Germany; Bernstein Center for Computational Neuroscience Berlin, Berlin, Germany; Einstein Center for Neurosciences Berlin, Berlin, Germany

**Keywords:** full-band EEG, DC-EEG, Cortical Spreading Depolarization, Brain Concussion, Mild Traumatic Brain Injury, Finite Element Analysis, Numerical Simulation, Electroencephalography, Electrocorticography

## Abstract

**Background:** Cortical Spreading Depolarization (SD) is a propagating depolarization wave of neurons and glial cells in the cerebral gray matter. SD occurs in all forms of severe acute brain injury as documented using invasive detection methods. Based on many experimental studies of mechanical brain deformation and concussion, the occurrence of SDs in human concussion has often been hypothesized. However, this hypothesis cannot be confirmed in humans as SDs can only be detected with invasive detection methods that would require either a craniotomy or a burr hole to be performed on athletes. Typical electroencephalography (EEG) electrodes, placed on the scalp, can detect the possible presence of SD but have not been able to accurately and reliably identify SDs.

**Methods:** To explore the possibility of a non-invasive method to resolve this hurdle, we developed a finite element numerical model that simulates scalp voltage changes that are induced by a brain-surface SD. We then compared our simulation results with retrospectively evaluated data in aneurysmal subarachnoid hemorrhage (aSAH) patients from Drenckhahn et al. (Brain 135:853, 2012).

**Results:** The ratio of peak scalp to simulated peak cortical voltage, Vscalp/Vcortex, was 0.0735, whereas the ratio from the retrospectively evaluated data was 0.0316 (0.0221, 0.0527) [median (1^st^ quartile, 3^rd^ quartile), n = 161, p < 0.001, one sample Wilcoxon signed rank test]. These differing values provide validation because their differences can be attributed to differences in shape between concussive- and aSAH-SDs, as well as the inherent limitations in human study voltage measurements. This simulated scalp surface potential was used to design a virtual scalp detection array. Error analysis and visual reconstruction showed that 1 cm is the optimal electrode spacing to visually identify the propagating scalp voltage from a cortical SD. Electrode spacings of 2 cm and above produce distorted images and high errors in the reconstructed image.

**Conclusion:** Our analysis suggests that concussive (and other) SDs can be detected from the scalp, which could confirm SD occurrence in human concussion, provide concussion diagnosis based on an underlying physiological mechanism, and lead to non-invasive SD detection in the setting of severe acute brain injury.

## 1 Introduction

Concussion, also known as mild traumatic brain injury (not to be confused with severe traumatic brain injury), affects 1.6-3.8 million individuals per year in the United States [1-3], and the actual number is likely higher because of underreporting and misdiagnosis [4]. In particular, sports concussion has garnered major public attention in recent years [5-9]. Concussions during military [10-12], automotive, occupational, and bicycle accidents have increasingly added to the public consciousness. Diagnosis of concussion depends on highly variable and often non-specific symptoms that include physical, cognitive, behavioral, emotional, and sleep-related aspects [13]. Immediate diagnostic methods rely on psychometric, reaction time, and balance assessments, but the accuracy, reliability, and repeatability of these assessments have come into question [14-21]. Health professionals typically test for potential sports-related concussions during the acute phase using the Sport Concussion Assessment Tool 5th Edition, SCAT 5, [22], which includes many of the aforementioned components. The assessment of acute concussion to prevent serious consequences of inappropriate return-to-play decisions [23, 24] is a well-established process that is based on multiple symptoms but without an underlying pathophysiological mechanism.

The historical [25] and emerging experimental [26, 27] evidence that concussion is accompanied by cortical Spreading Depolarization (SD) suggest that SD might be the underlying cause of, or at the least contribute to, acute concussion symptomatology in the immediate minutes after concussion. Currently, diffuse axonal injury (DAI) is the most widely accepted anatomical mechanism accounting for the acute and chronic symptoms of concussion. DAI has been proposed as an objective diagnostic criteria based on axonal pathophysiology assessed by MRI and the increased blood concentration of axonal proteins [28]. However, the effects of DAI and SD are separated both by where they occur anatomically, with DAI affecting axons and SD occurring in neurons and glia, and the timing of their effects. Concussion is thought to induce a brain network dysfunction, in which white matter damage and DAI are implicated in characteristic symptoms such as slowed information processing at one day [29] and five years [30] post-concussion, time periods which are not relevant to the immediate occurrence of SD following concussion.

### 1.1 What are SDs?

SDs are regions (typically 3 mm width or diameter [31, 32]) of large amplitude direct-current (DC, 5 - 30 mV) negative *en mass* depolarization of neuronal and glial cells in gray matter that persist over several minutes [33-37]. The hallmark of SD is the near-complete breakdown of the neuronal transmembrane ion gradients [38] that leads to influx of water and cytotoxic edema [39]. Secondarily, glutamate is released in large amounts [40]. SDs are slow-moving waves (1 - 9 mm/min) that lead to depression of electroencephalographic (EEG) amplitudes for 0.5 - 3 minutes, with normal function grossly restored after 5 - 10 minutes if the vasculature is not compromised [31, 41, 42]. Although SDs were originally discovered in rabbits in the 1940s [31], it was not until the 1990s that they were initially observed in a single severe acute brain trauma patient [43]. In the early 2000s, Strong and colleagues presented the first robust bedside method that used subdural electrodes to detect SDs in approximately 50% of individuals with traumatic brain injury [36].

SDs occur abundantly, periodically, and repetitively in severe acute brain injury, including malignant hemispheric stroke, subarachnoid hemorrhage (SAH), spontaneous intracerebral hemorrhage, and severe traumatic brain injury [41, 44-49]. The transition from long-lasting SDs in severely ischemic tissue to short-lasting SDs in less ischemic or normal tissue has been described as “the SD continuum” [50, 51]. Terminal SDs during the development of stroke, brain death and death from circulatory arrest represent the extreme end of the SD continuum [34, 52-58]. In addition, short-lasting SD in otherwise healthy tissue is assumed to be the underlying cause of migraine aura [35, 59, 60], which is the only clinical manifestation of SD in which the wave-like nature of SD becomes clinically apparent, leading to the fallacy for many decades that migraine aura is the only clinical manifestation of SD [50]. In the first phase of ischemic strokes due to arterial occlusion, and most likely in concussion, the SD spreads concentrically from the ischemic core [61] or the point of impact [31]. The consequences of SD occurrence in both migraine and concussion, if it does occur in human concussion, are temporary, in stark contrast to the “continuum” pathological role of SD in severe acute brain injury [50].

### 1.2 Rationale for SD Occurring in Concussion

SD and concussion were first linked by observation that a firm, non-damaging touch of the brain elucidated an SD [31]. This initial observation of brain deformation causing SD was the basis for extensive studies using controlled brain deformation [62-64]. These studies of brain deformation were not focused on human concussion but rather the mechanical parameters of brain deformation, including the speed, area, and depth of the depression that elicited SD. The direct link between concussion and SD was suggested based on a comparison of the energies required to elicit SD and experimental concussion [65]. This comparison is weakened by concern that: 1) the calculation of energies was not described or referenced; 2) only experimental, non-human studies provided evidence; and 3) the rate of deformation, a parameter that determines SD initiation, was not used. These concerns make the presumption that human concussion is always accompanied by SD a hypothesis that needs testing.

Many experimental studies have noted the similar features of concussion and SD, including high metabolic rate [66] and high [K^+^]_o_ [67]. In particular, EEG suppression [68-70] and negative DC-potential [70] in experimental models of concussion were consistent with the original observations of EEG suppression [31] and negative DC-potential during SD [37].

Studies demonstrating the presence of SD in mouse models of concussion strongly suggest that SD might be a common feature of concussion [26, 27]. The behavioral effects associated with SD after controlled cortical impact without craniotomy are similar to the diverse symptoms used to diagnose human concussion [26]. All of these experimental studies used models of concussion that excluded cerebral contusion or intra-cranial hematomas.

Several clinical studies [71, 72] and reviews [73-75] have presented compelling arguments suggesting that SD could explain concussion symptomatology. In particular, Oka et al. [71] reported that patients with non-convulsive concussion-like injury showed symptoms of “headache, nausea and vomiting, pale complexion, somnolence, irritability or restless confusion” that are “also characteristic symptoms of classical migraine attacks.” Although this experimental and clinical evidence that a blow to the cranium could trigger SD is scientifically valid, the mechanism of SD initiation by brain deformation has not been explored and the electrophysiological confirmation of this scenario in humans [25; page 1083] is missing.

### 1.3 Mechanisms of Mechanical Brain Deformation Induced SD

Although the mechanisms of SD initiation and propagation have been extensively explored [25, 76], there has been no attention devoted to the mechanism of SD elicited by brain deformation. Strangely, brain deformation has been ascribed to many modes of eliciting SD, as thoroughly described by Marshall [77], including pinprick, pledget application of compounds that elicit SD to the cortical surface, neurosurgical procedures that involve brain deformation or compression, and others. However, many studies have been performed using *in vitro* models of neuronal stretch that show both permeability and ion channel changes [78, 79] whose time scale is relevant to concussive SD initiation. In this same time scale, there is evidence that cells do partially depolarize after stretch due to permeability increases [80, 81]. These issues could be explored for their connection to the mechanism of SD generation by brain deformation. A recent study suggests spontaneous SDs in an “intracortical hemorrhage model are triggered by the mechanical distortion of tissue by rapidly growing hematomas” [82].

To explore the possibility of non-invasive detection methods that might resolve the conundrum between the scientific evidence for, and missing confirmation of, SD in human concussion, we performed a numerical simulation to model the spatial and temporal characteristics, as well as the magnitude, of the scalp DC-voltage from an assumed brain-surface concussive SD. We then compared the computational model with retrospectively evaluated human results from aneurysmal SAH (aSAH) patients, given that human concussion data are not available. Finally, we used this simulated scalp DC-potential to investigate the ideal spatial separation of the electrodes in a sensor array for scalp DC-EEG detection of SD.

Our hypotheses are that our simulated scalp DC-potential: 1) compares with existing experimental data; 2) can be used to determine the electrode configuration, specifically the electrode spacing, that enables, and is sufficient for, the accurate reconstruction of a scalp DC potential image that aids in the visual detection of SD; and 3) is consistent with the detection limits of DC-EEG. The implications of these hypotheses are that if propagating SDs can be noninvasively detected, concussion detection using SD is possible, the role of SD in concussion could be explored, and noninvasive SD detection in severe acute brain injury patients might be possible.

## 2 Methods

### 2.1 DC-EEG Forward Problem using a Finite Element Model

Various authors have conducted numerical simulations of EEG phenomena in the brain and head directed both at the forward problem [83] and inverse source localization of the focus of epileptic seizures [84, 85] and of regions of suppressed neural activity [86]. Most models focused on the number of tissue layers and geometry [83, 84, 87-90] but others have been concerned with material properties and constitutive equations [91-93]. However, all these models used clinical EEG frequencies greater than 0.5 Hz, except for the inverse source localization effort by Miller et al. [85] which used frequencies of 0 to 70 Hz in contrast to the clinical frequency range. Our forward problem simulation is unique as it deals with the near-zero frequencies of an SD that can be modeled with just a brain-surface DC-potential without any underlying cortical structure and depends on DC-EEG, not the normal clinical EEG frequencies above 0.5 Hz.

#### 2.1.1 SD modeled as a brain surface voltage field

To implement our simulation, we first mathematically describe the SD voltage waveform. The cortex was not modeled as the SD can be described as an electric field on the cortical surface. The SD voltage waveform on the brain-surface is instead implemented into the model with the boundary condition:

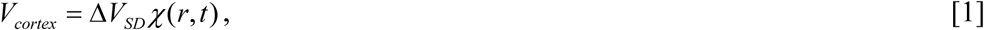

where ΔV_SD_ is the cortical surface voltage potential of SD (−20 mV) and *χ* is a shape function describing the spatio-temporal behavior of the spreading, namely an expanding ring with a thickness or width of 3 mm [31, 32] and a velocity of 3 mm/min appearing within 1 min after impact [26, 27].

Two different shape functions were used to describe the SD, whose initial inner radius is zero at the concussive focus. The first shape function is a square wave:

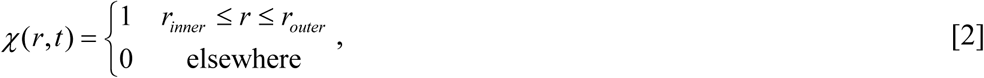

where r is the distance from the focus of the concussion. The radius of the ring expands outward as:

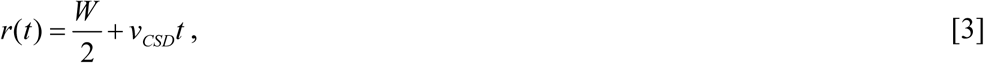

where W is the constant ring thickness of the wave (3 mm), and v_SD_ is the propagation speed of the SD (3 mm/min). The inner (trailing) edge of the ring, r_inner_, and outer (leading) edge of the ring, r_outer_, are then

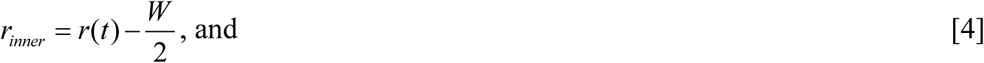

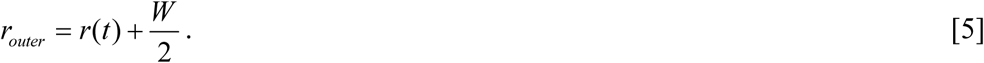

The second shape function is a triangular wave:

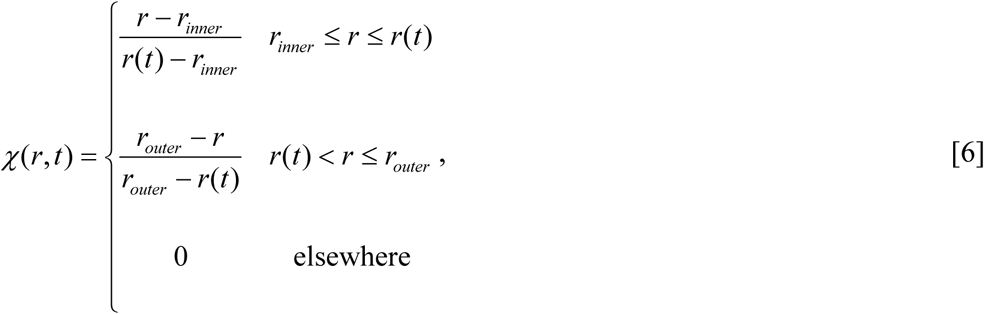

where r(t) is the radius of the expanding ring, and r_inner_ and r_outer_ are again the inner and outer radii of the ring. These two shapes bracket the largest and smallest transmission of signal to the scalp (or most optimistic and least optimistic signal estimates), including the more typical shape observed using ECoG methods and the shape of an SD DC shift in an experimental model of concussion [26], for a given SD voltage, ΔV_SD_, and width, W. These boundary conditions (Equations 1-6) define the spatial extent of the modeled SD that are needed to determine the solutions to Equation 7 below.

#### 2.1.2 Layers between the brain surface and scalp through which the SD electric field must pass

Our next step was to model the layers between the brain surface and the scalp through which these brain surface voltage fields must travel to reach the scalp. A five-layer model was used for this project, focusing on 1) cerebrospinal fluid (CSF); 2) dura matter; 3) skull; 4) muscle; and 5) skin (see Fig. 1b). Ramon et al. [84] showed that the CSF was the major factor in a quality model. Muscle was included to account for the portions of the frontal bellies of the occipitofrontalis muscle that attach to the *galea aponeurotica* [94]. Although the muscle layer is thin, it was included for completeness and adds to the truthfulness of the model. The total thickness is 1.02 cm and the thickness of each layer can be found in Table 1. The values for layer thickness, relative permittivity (dielectric constant), and conductance, as presented in Table 1, were obtained from various literature sources [83, 84, 93, 95, 96]. The frontal bone of the skull was chosen as the reference because this is the area where athletic concussions are most likely to occur, and it is also the thickest part of the skull on average, resulting in the greatest attenuation of the signal [97]. Furthermore, the results will be compared to clinical research data collected by Drenckhahn et al. [34] for SDs occurring primarily in the frontal cortex of aSAH patients (see below for details). Instead of relying on a multilayer model of the skull, i.e., hard and soft bone [83, 98-100], the theory of interacting continua was used to determine the parameters through mixture theory [101].

**Fig. 1.**
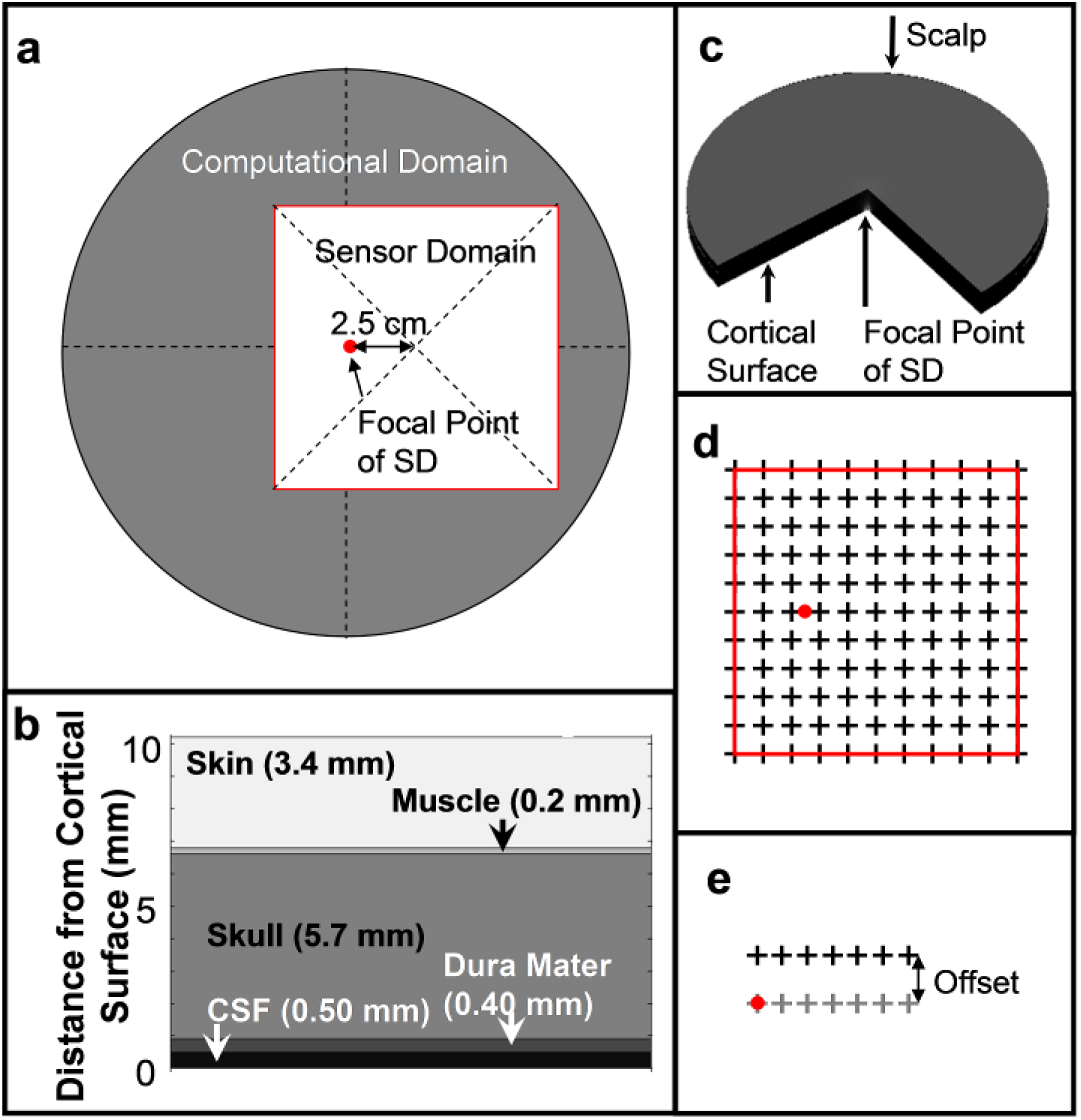
The associated geometry and locations for the numerical analysis. **a**) The overlay of the 20-cm diameter computational domain (dark gray) and the sensor domain (white square with red border) showing the focal point of the spreading depolarization (SD) as a red dot. **b**) A slice view of the model layers with thickness representing cerebrospinal fluid (CSF), Dura Mater, Skull, Muscle, and Skin. **c**) The three-dimensional computational domain with a 90° quadrant removed to show the layers and focal point of the SD. **d**) An example of the 10 cm x 10 cm sensor grid (red border) with 1-cm spacing and electrode locations indicated by crosses and the focal point of the SD as a red dot. **e**) The sensor offset for velocity and geometric calculations with the ideal sensor position with gray crosses, an offset array in black, and the focal point of the SD as a red dot. The offsets used were 0.1, 0.5, and 1.0 cm

#### 2.1.3 Calculation of the SD electric field propagating from the brain surface to the scalp

Our further step was to calculate the propagation of the electric field through these layers from the brain surface to the scalp using Poisson’s equation:

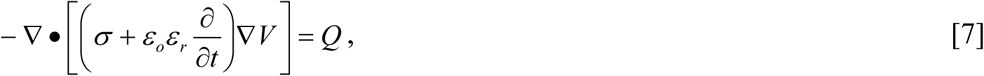

where t is time, σ is the conductivity, ε_o_ is the permittivity in a vacuum, ε_r_ is the relative permittivity, V is voltage potential, and Q is a source term. The boundary condition at the scalp was zero normal current density or Q = 0. No other current sources (Q) were used; however, Q could be used to produce noise and additional signal generation such as that of the transdermal epithelial potential (TEP) or the galvanic skin potential. The model ignores the TEP, i.e., assumes the skin is abraded [102, 103]. In more detail, Poisson’s equation relates the contributions from the voltage field on the cortical surface, *V*_*cortex*_, defined by the boundary condition (Equation 1) and its associated shape functions (Equations 2 and 6), as it propagates through the various layers from the brain surface to the scalp. The calculation was implemented using finite element modeling, a method that creates a mesh of cells between which these equations are solved numerically. These equations of electrodynamic theory and their use in finite element modeling are inherent in computational neuroscience tools such as FieldTrip [104, 105] that are used to study the forward and inverse problems. These tools are focused on clinical EEG frequencies in contrast to our focus on SDs DC potential.

### 2.2 Simulation Details and Parameters

The simulation was performed on a cylindrical domain with a radius of 10 cm (see Fig. 1a and 1c). This radius was chosen to be large to prevent interference effects of the boundary condition. Due to the slow propagation velocity of SDs, the total time simulated was 15 minutes as a pilot study showed little change in the basic structure of the propagation after this time. The simulation was performed using COMSOL 4.3 (COMSOL, Inc.) with a structured hexagonal mesh, which uses an adaptive time solver with a relative error below 1e-6. The spatial resolution of the mesh was evaluated using the grid convergence index (GCI) method, which is the American Society of Mechanical Engineers standard [106]. The mesh element sizes were 0.886, 0.481, and 0.246 mm for the coarse, medium, and fine meshes, respectively. The coarse mesh had some elements with GCI qualities as low as 0.4 (<10% of elements) and the fine mesh elements all had quality values above 0.95; all elements had GCI qualities above the level of concern (0.1). Furthermore, the simulation was validated against an analytical solution for a constant voltage drop of 20 mV applied between the scalp and the cortex. The layer resistances, R, can be calculated as:

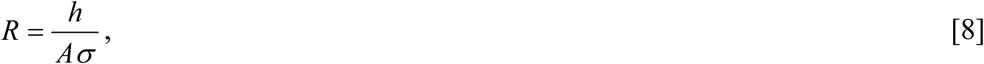

where h is the thickness and A the cross-sectional area of the layer. The current and voltage drop can then be calculated from Ohm’s law. The resulting errors were less than 3e-7 mA and 1e-5 mV for current and voltage, respectively, between the closed-form solution and the numerical results.

The shape of the potential of the SD as it propagates through the various layers was characterized through a relative voltage, V_rel_:

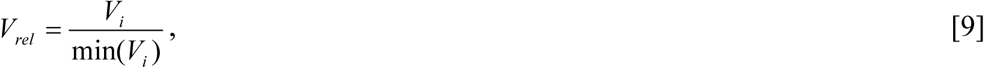

which compares the voltage of the i^th^ layer, V_i_, to the minimum voltage on that layer occurring between the time of 5 and 15 minutes, min(V_i_). A value of 0.2 was set for the threshold of V_rel_ in order to determine the thickness or width of the SD-based DC-potential and to tag times for analyzing sensor data.

### 2.3 Scalp Potential Image Reconstruction, Simulated Scalp Detector Construction, and Reconstruction Error

This simulation/modeling process of an SD’s DC electric field on the brain surface that transverses the various tissue layers from the brain to the scalp results in an estimate of the electric field on the scalp. This electric field is then “sampled” by an array of electrode sensors at various spacings. The final step is then the reconstruction of the scalp electric field from this electrode data so that the scalp electric field can be presented as a heat-map image. The errors between the known scalp electric field and its reconstruction from the electrode sensors can then be evaluated. Errors in velocity were determined in a subsequent study using the scalp electric field calculated as a results of the simulation, but with a limited electrode array.

The scalp surface signal over a 10 cm x 10 cm square (Figure 1a) was reconstructed from simulated electrode readings by sampling the predicted voltages at various electrode spacings. The simulated SD was centered at 2.5 cm from the left of the center of the sensor array as shown in Figure 1d. The relative location of the sensor grid to the simulation or sensor domain and focal point of the SD can be seen in Figure 1a. Figure 1d shows a sample grid with 1-cm spacing. Reconstruction of the scalp electric field was performed with bilinear interpolation between four electrodes [107]. The root-mean-squared error (or the L2 norm),

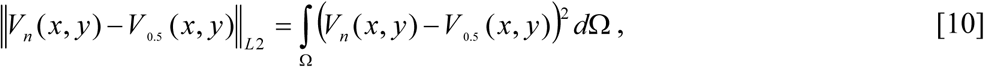

where Ω is the domain, was used to compare the error between the voltage reconstruction at 0.5-mm electrode spacing (V_0.5_) and the other arrays with electrode spacing of 1, 5, 10, 15, 20, 25, 30, 40, and 50 mm (V_n_). The reconstruction was also performed by averaging the voltage potential over circular areas (0.5, 0.75, 1, 1.5, 2, 2.5, 3, 4, and 5 mm radii) representing various sizes of the active area of the electrode. Results were compared with the infinitesimal or point values of voltage.

### 2.4 Errors in Velocity and Ring Thickness Evaluated using Detection Array Offsets

A second study, independent of the regular grid study above, was performed to understand how different offsets (see Fig. 1e) influenced SD velocity calculations from the simulated scalp data. To simplify this error analysis, a subset of the full electrode array was used composed of a linear electrode array. Errors in the velocity of the SD will occur if a linear sensor array is not aligned with the velocity field, which occurs for a concentrically expanding wave when the sensors are offset from a given radius. Therefore, velocity and ring thickness calculations were performed for a 1 × 7 array of electrodes with the sensor array situated so that the first electrode is located over the focus of the SD, and then with 3 additional linear 2 × 7 arrays that were separated by 0.1, 0.5, and 1 cm. All arrays had an electrode spacing of 0.5 cm and length of 3 cm (see Fig. 1e). An apparent velocity, v_app_, was calculated using the formula:

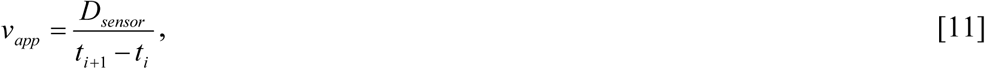

where D_sensor_ is the spacing between two electrodes and t_i_ is the time at which V_rel_ of the i^th^ electrode either first exceeds (onset) or is first restored below the threshold of 0.2 (abatement). Velocities were calculated using each electrode pair (i and i+1), with the first pair consisting of the two electrodes closest to the focus and so forth. A scalp depolarization time (t_d_), where the threshold value was exceeded, was defined to be the time between onset and abatement. The scalp ring thickness (W_s_) was calculated using the equation:

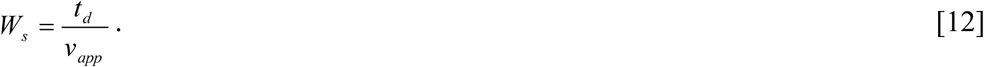

For this second study, the scalp DC potential of a brain surface expanding ring SD was simulated with a velocity of 3 mm/min, a ring thickness of 30.8 mm, and a depolarization time of 10.3 minutes, values derived from the FEM simulation and the time needed to transverse the length of the simulated linear electrode array.

### 2.5 Retrospective Evaluation of Human Data [34]

We retrospectively evaluated the amplitude of SDs in three aSAH patients with sufficient quality subdural full-band ECoG (fbECoG) and scalp full-band EEG (fbEEG) recordings from Drenckhahn et al. [34]. Patient recruitment and characteristics, fbEEG and fbECoG recording, data processing and analysis are detailed in Drenckhahn et al. [34]. Research protocols were approved by the institutional review board and surrogate informed consent was obtained for all patients. All research was conducted in accordance with the Declaration of Helsinki.

In brief, a single, linear, six-contact (platinum) recording strip (Wyler, 5 mm diameter; Ad-Tech Medical) was placed on the surface of the cortex for ECoG recordings [36, 44, 45]. Both fbECoG and fbEEG recordings were performed (bandpass: 0–1000 Hz, sampling rate: 2500 Hz) using a BrainAmp amplifier. The electrodes were attached with Collodion adhesive (Mavidon) after abrasive electrode gel (Abralyt 2000, EasyCap) and conductive electrode cream (Synapse, Med-Tek) were applied to minimize electrode impedance (<5 kV) and ensure long-term stability of the signal. Data were recorded and reviewed with the use of LabChart 7 software (ADInstruments) and BrainVision Recorder 1.05 software (Brain Products), respectively. For each pair of subdural fbECoG and scalp fbEEG recordings from the same SD, the maximum voltage of each was recorded. The ratio of these subdural and scalp voltages was compared with the ratio determined from our simulation.

### 2.6 Statistics

Statistical tests were performed using SAS/STAT version 15.1 (SAS institute Inc, Cary, NC). A nonparametric statistical test was used because the voltage ratio measurements deviated significantly from a normal distribution. The one sample Wilcoxon signed rank test was used to compare our simulation derived voltage ratio with the voltage ratio determined from data retrospectively extracted from the work of Drenckhahn et al. [34]. Data are reported as medians (1^st^ quartile, 3^rd^ quartile) and p < 0.05 was accepted as statistically significant.

## 3 Results

The simulation results for the voltage and current density distributions in the various layers from the cortex to the scalp for a cortical triangular wave after 5 seconds and 10 minutes are shown in Fig. 2. The voltage field and ring shape at the interface between each layer of the model starting at the cortex and continuing to the scalp are summarized in Table 2. Early in the migration of the voltage depression, the voltage requires more time to return to rest after passage of the wave peak when compared to later time points. On the scalp, the relative voltage exceeded the 0.2 threshold at 14.6 mm ahead of the peak voltage and remained elevated 16.2 mm behind the peak value. Figures 2b and 2d show that the largest current densities occur in the first layer composed of CSF, as it is the most conductive, peaking at 35.6 mA/m^2^ at the point of peak voltage on the cortical surface and weakening to below 2.1 mA/m^2^ at locations 2 mm before and after the value of SD radius, r(t), at the peak cortical voltage. The muscle tissue also shows an elevated current density (20 mA/m^2^), though much smaller than that in the CSF.

**Fig. 2.**
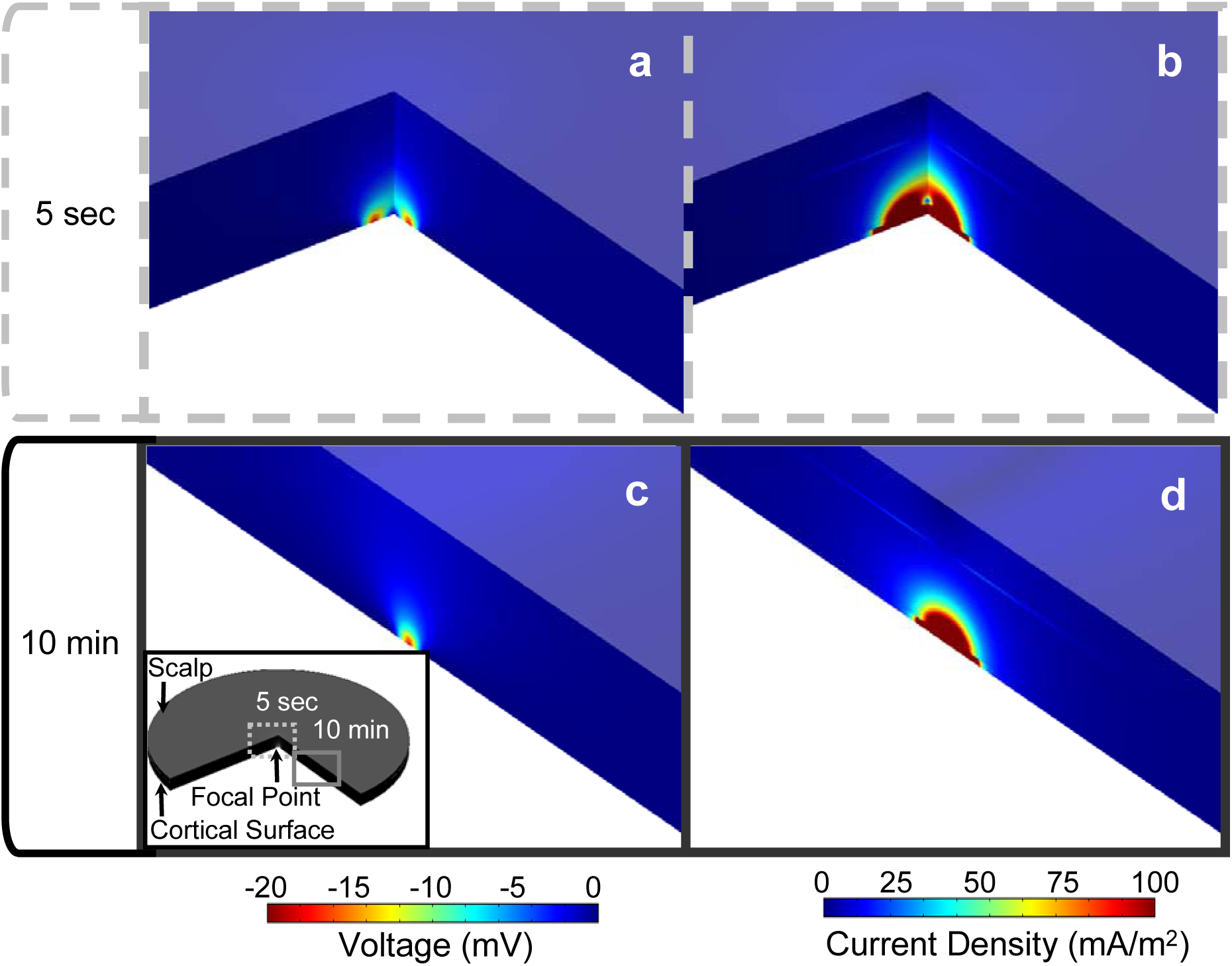
Simulation results for electrical voltage and current density at 5 sec and 10 min after spreading depolarization (SD) initiation in the 10-cm disk. Panels **a** and **b (**dashed border) show the numerical results at 5 seconds, and panels **c** and **d (**solid border) show them at 10 minutes after initiation of the SD for a concentrically spreading SD propagating radially. Panels **a** and **c** show the electrical potential (voltage in mV), and panels **c** and **d** show the current density (mA/m^2^). The numerical pseudo-color scales for voltage and current density are shown below panels **a** and **c**, and panels **b** and **d**, respectively. The insert in panel **c** shows the relative location of the views of the SD depicted in panels **a** and **b** (5 seconds) as well as **c** and **d** (10 min). Also included in the insert are the location of the scalp, cortex, and focal point of the SD

### 3.1 Voltage Decrease

The voltage detectable at the scalp is attenuated to 0.0735 of its strength at the cortex as shown in Table 2. The voltage is attenuated through each layer, while the affected region grows larger. The largest amplitude drop, ΔV = 8.18 mV, occurs in the skull. A drop of 5.10 and 5.01 mV occurred in the CSF and dura matter, respectively. The total voltage amplitude drop across the skin and muscle taken together was only 0.240 mV. Figure 3 compares the voltage at the cortex to that of the scalp, and at each interface between layers.

**Fig. 3.**
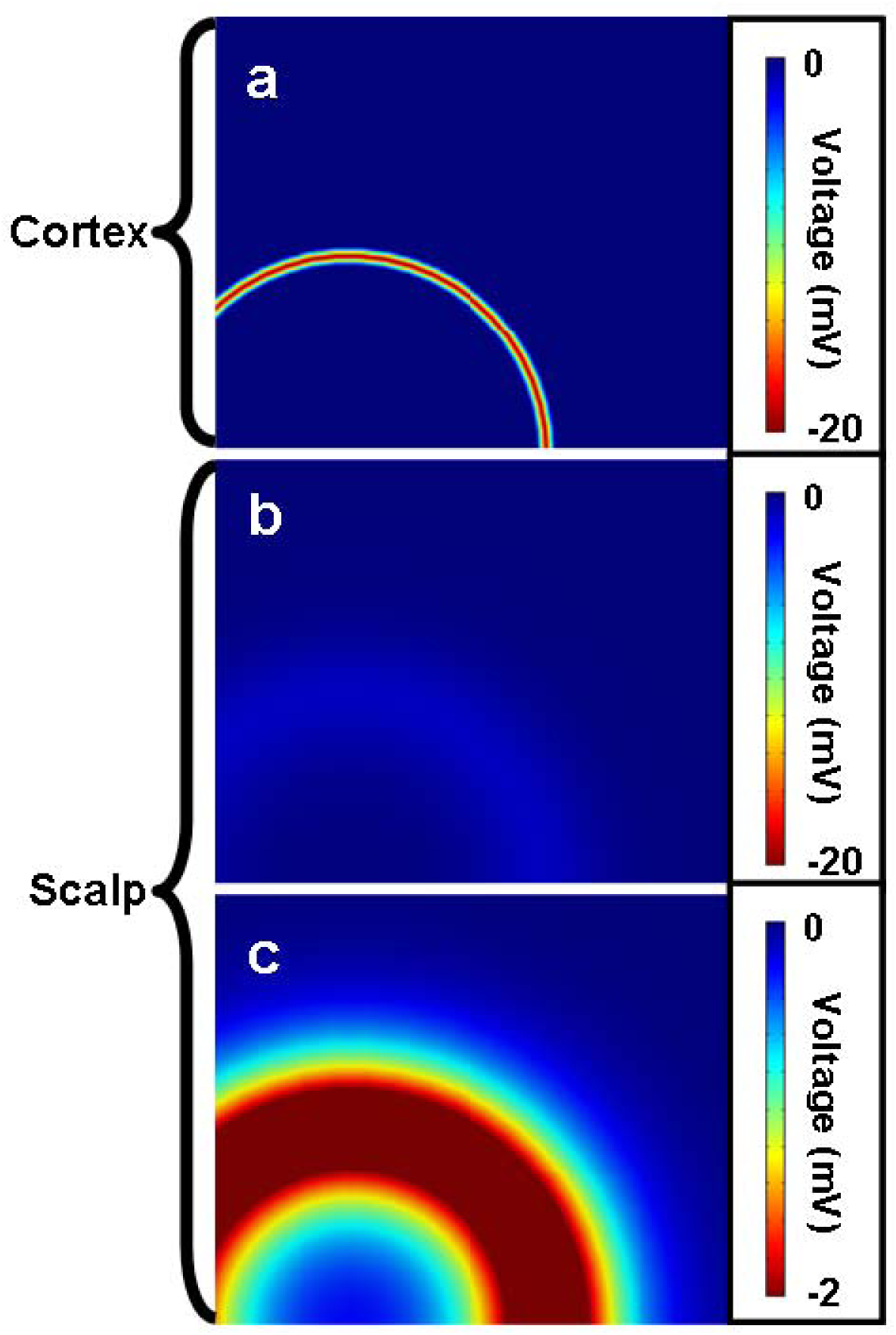
The simulated voltage potential of a cortical spreading depolarization diminishes in amplitude at the scalp (**b**) but increases in effective width (**c**). The voltages in the cortex and scalp are shown with identical scales in (**a)** and (**b)** to highlight the change in amplitude. The scalp voltage map is rescaled in **c** to highlight the increase in ring diameter

### 3.2 Dimensional Changes

The area above the threshold voltage grew most in the skull (595%) and least in the muscle (0.74%). The thickness of the voltage ring was 2.4 mm at the cortical surface and increases to 30.8 mm (x 12.8) at the scalp. There is also a spatial lag between the locations of the peak voltage on the cortex and the scalp of 0.474 mm, which corresponds to a time delay of 0.16 minutes. Supplemental Movie S1 presents a movie of the scalp voltage from a cortical SD with both top and side views.

### 3.3 Voltage Ratios from the Simulated Square and Triangle Waves

This study investigated two types of propagation waves: a square wave and a triangular wave. Their peak voltages from the scalp electrode sampling results were noticeably different: V_peak_ was 3.6 and 1.47 mV for the square and triangle waves, respectively, giving relative voltages of 0.18 and 0.0735.

In addition to simulating the scalp signal with tissues having electromagnetic properties as measured in Penn and Bell [95], we also simulated the triangle wave profile with tissue having properties as reported by Gabriel et al. [96]. Though the estimates of these parameters were different, some by several orders of magnitude, the resulting peak scalp voltage ratio was 0.09, as opposed to 0.0735, suggesting that signal detectability at the scalp is not sensitive to the variation of tissue properties between (or within) subjects.

### 3.4 Voltage Ratio Comparison with Clinical Research Results

We compared the voltage ratio from our simulation results with the voltage ratio retrospectively evaluated from the data collected by Drenckhahn et al. [34]. Paired and identifiable fbECoG subdural and fbEEG scalp recordings were available for 161 SDs. In the pooled analysis, the maximum fbECoG subdural peak amplitude among all electrodes for each individual SD was -9.3 (−7.2, -10.1) mV. In the scalp fbEEG recordings of the same SDs, the maximum peak amplitude among all electrodes was -273 (−376, -176) µV. The ratio of peak scalp to peak subdural voltage was 0.0316 (0.0221, 0.0527) in the pooled data analysis (n=161). The simulation results differed from the retrospectively evaluated data; the ratio of peak scalp to simulated peak cortical voltage, Vscalp/Vcortex = 0.0735 versus 0.0316 (0.0221, 0.0527), n = 161, p < 0.001, one sample Wilcoxon signed rank test.

### 3.5 Effect of Electrode Spacing on Visualization of Scalp Voltage

Figure 4 shows the reconstruction of the surface signal using regularly spaced sensor arrays with varying separation distance. The voltage reconstructions are visually consistent with the true field up to a sensor grid spacing of 1.0 cm. At larger electrode grid spacings of 2 cm and above, a breakdown of the ring structure occurs, and at even greater spacings the structure could not be discerned. Figure 5 illustrates how the loss of structure occurs due to the change in grid spacing starting at the 2 cm spacing (Figure 5c) and continuing at the 3 cm spacing (Figure 5d). Supplemental Fig. S1 shows the dependence of the degradation of the true scalp voltage on the electrode spacing and the time after SD initiation.

**Fig. 4.**
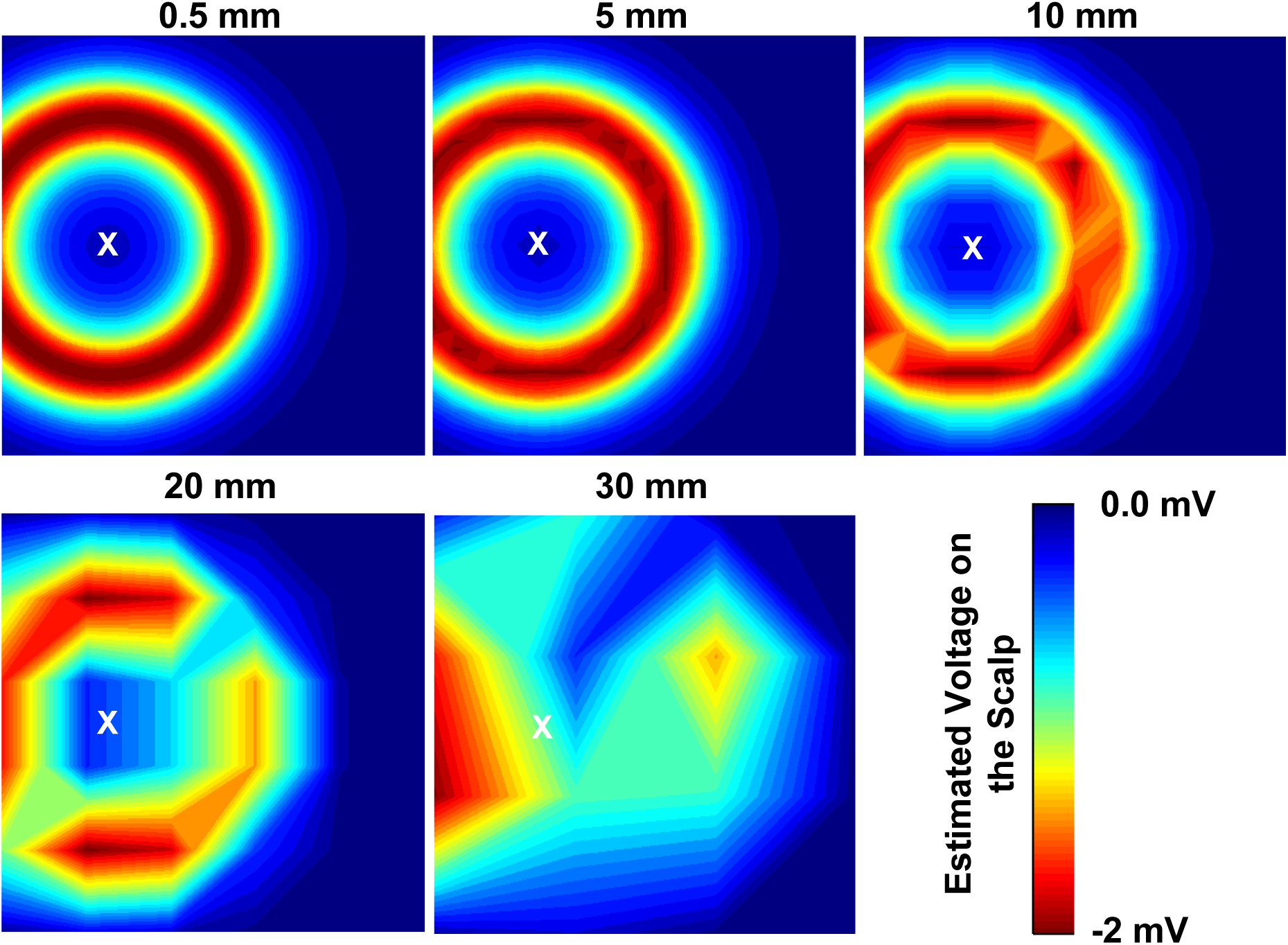
The scalp voltage was sampled and reconstructed using simulated sensor grids with electrode spacings of 0.5, 5, 10, 20, and 30 mm. These voltage maps represent the signal at 10 minutes after the onset of the spreading depolarization (SD). They illustrate the degradation of the reconstruction with increasing electrode spacing, and the preservation of the scalp voltage at electrode spacings of less than 1 cm. The white X indicates the focal point of the SD. Reconstruction was performed using bilinear interpolation between four electrodes

**Fig. 5.**
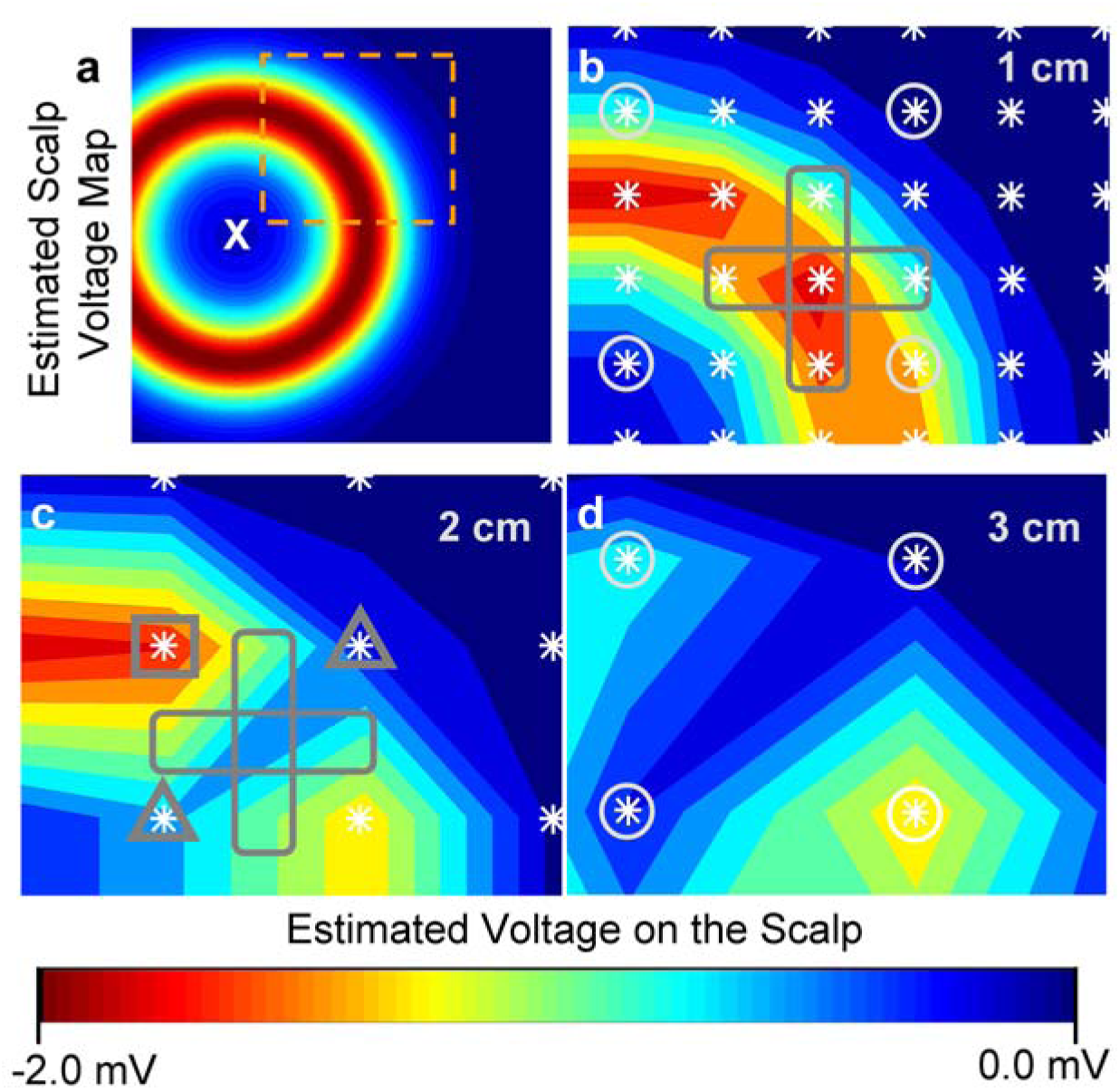
Numerical simulation results of the scalp voltage reconstruction of a spreading depolarization (SD) expanding ring with different electrode spacings. (**a**) Estimated scalp voltage intensity map at 10 min after SD initiation in a 10 cm x 10 cm simulation domain. The symbol X marks the focal point of the SD on the underlying brain surface. The region within the dashed orange box is expanded to show the detail of the detected signal degradation with increased electrode spacing in (**b)**, (**c)**, and (**d)**, with electrode spacings of 1, 2, and 3 cm, respectively. (**b**) Only the 1 cm spacing shows the full ring structure of the scalp voltage (∼2.5% error). (**c**) The 2-cm sensor spacing shows a recognizable ring but a low voltage wedge has broken the structure (∼15% error). The dark cross (**b** and **c**) indicates the locations of sensors that are removed from the 1 cm design to obtain the 2-cm design, which result in the breakdown of the ring. With the removal of the signal from the electrode sensors within this cross, the reconstructed heat map image losses data points that contribute to the ring structure. (**d**) The 3-cm spacing fails to reconstruct the ring structure in the intensity map (∼25% error). The lost of the signal from the electrode marked with a surrounding square, located on the ring of the SD (depressed voltage), and from the two electrodes marked with surrounding triangles, on the normal tissue (baseline voltage), results in the development of a notch, thus breaking the ring structure. The white circles (**b** and **d**) indicate the sensors from the 1-cm design that remain in the 3-cm array

### 3.6 Effect of Electrode Spacing on the Error in the Reconstruction of Scalp Voltage

Figure 6 shows the root-mean-squared error of the reconstructed scalp voltage as a function of electrode spacing: the error approaches 5% for electrode spacing of 1.5 cm and increases to 25% for electrode spacing above 2.5 cm. When taking into account differences in possible electrode diameter from 0.1 to 1 cm, we found that the reconstructions only varied by 2% even when comparing a point electrode to one that was 1 cm in diameter. The error in reconstructing the brain-surface field from an electrode grid could be further reduced by application of more complex and computationally expensive interpolation methods.

**Fig. 6.**
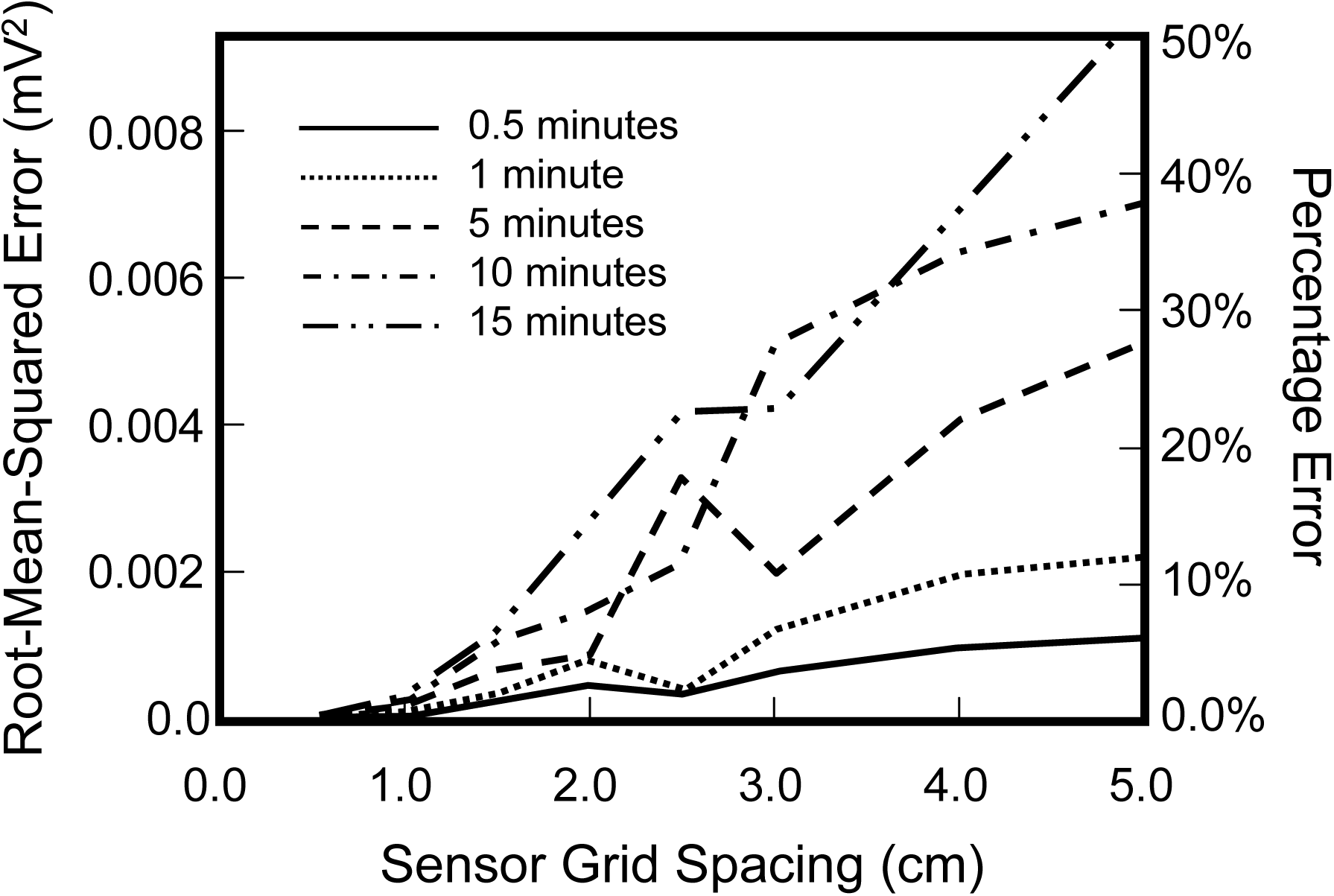
The root-mean-squared error in the reconstruction, using the 0.05-cm spacing as the baseline, for each of the sensor designs as a function of sensor electrode spacing and time. The error is reported at 0.5, 1, 5, 10, and 15 minutes

### 3.7 Errors in Velocity and Ring Thickness

The simulation results allowed the velocity, as one of the characteristic parameters of SD, to be estimated from the simulated scalp potential, but as previously noted, errors were dependent on a misalignment of the strip electrode and the origin and path of the spreading wave. The velocity and ring thickness predictions were dependent on the sensor offset from the velocity vector, as well as on the proximity to the focus of the SD, and reached values less than 10% and 4% for velocity and thickness, respectively.

The velocity was consistently over predicted. The error in the velocity calculation (in relation to the simulation velocity of 3 mm/min) using the onset time (V_rel_ exceeding 0.2) was highest near the focal point of the concussion, being 127%, 127%, 117%, and 284% for the 4 different offsets, and decreasing to 7.5%, 6.4%, 8.7%, and 19% for the final electrode pairing, i.e., the two electrodes furthest from the focus of the SD. The errors in the velocity calculation were smaller using the abatement times (V_rel_ returns to less than 0.2) with two electrodes closest to the focus predicting errors of 82%, 82%, 88.7%, and 122%, but the velocity errors were similar at the final electrode pairing to those predicted using the onset time.

The average time of electrode activation for V_rel_ > 0.2 was 9.66 (9.12, 9.9) minutes excluding electrodes for which the wave did not completely pass within the 15-minute time frame. The error in ring thickness increased as the array was offset; the error between predicted and actual thickness was 3.51% to 9.8% for the zero and 1 cm offset arrays, respectively.

## 4 Discussion

We simulated the scalp voltage produced by a concentrically traveling SD on the cortex of the brain and estimated that an electrode spacing of 1.0 cm is required to accurately collect the spatio-temporal data of the SD with image reconstruction errors of less than 5%, whereas an electrode spacing of 2 cm gives reconstruction errors greater the 15% at 15 min (see Figure 6). The results showed that the SD wave detectability is dependent on electrode grid spacing, with loss of structure as grid spacing increases. The ratio of simulated peak scalp to peak cortical voltage (Vscalp/Vcortex = 0.0735) was approximately double compared to the retrospectively evaluated clinical research observations of Drenckhahn et al. [34]. This difference is acceptable as its due to differences in SD shape and configuration between our simulated concussive and aSAH SDs. Although the amplitude from the simulated concussive SD is diminished to ∼7% of that at the brain surface through resistive dissipation, it is large enough to be detected by current DC-coupled amplifiers. The spatial expansion of the negative DC-potential at the scalp of a brain-surface SD, which occurs mostly in the skull (low current but high resistance), aids in the non-invasive detection of a SD as it enlarges the ring thickness at the scalp.

Our simulation results were highly accurate (peak errors < 1e-3%) when compared to the closed-form solutions. This accuracy suggests that imaging principles that were used in this work could be applied to detection and tracking of SDs. Therefore, systematic errors due to our model’s simplified geometry and assumed uniform thickness of the layers and to the effect of variations in material properties (conductivity and relative permittivity) used in our simulation will not detract from the accuracy and reliability of the simulated data for its potential use in future designs.

### 4.1 Existing Methods of SD Detection

The current method for detecting and measuring SDs using electrocorticography (ECoG) with strip electrodes directly placed on the cortical surface [36, 47] is invasive. This method is only used on patients where the electrodes are placed concurrently with a prior necessitated craniotomy or a burr hole trepanation for placement of a ventricular drain or oxygen sensor [108] as performed for the planning of resective epilepsy surgery [109; see section ‘Burr Holes for Placement of Strip Electrodes’ (4th page) and Figure 4]. Often, publications using ECoG SD detection highlight the lack of a scalp detection method to reliably detect SD non-invasively [35, 108].

### 4.2 Previous Non-Invasive Detection Attempts

To answer this need for non-invasive SD detection, there have been two efforts directed at scalp SD detection using simultaneous ECoG at the cortical surface and EEG at the scalp with clinically-based electrode positions [34, 110] the spacings of which are typically estimated as ∼8 cm for the 10-20 and ∼2 cm for the 10-5 system [111, 112]. Stationary residual negative scalp DC voltages between 65 μV and 1 mV associated with propagating SDs using ECoG have been measured on the scalp by Drenckhahn et al. [34]; however, SD propagation was not observed on the scalp (see Fig. 1A, traces 1-3 and 10-12, in Drenckhahn et al. [34]). The widened DC-potential at the scalp shown by our simulation is possibly the reason Drenckhahn et al. [34] observed that the scalp DC-voltage surface appeared at multiple electrodes simultaneously.

In another study, scalp observations of EEG suppression and DC-potential changes in patients whose bone-flap had been removed (except for 1 of 18 subjects) were guided by ECoG-detected SDs. However, unguided scalp SD detection was not established [110]. A replication of this study [113] and a follow-up rebuttal [114] did not provide evidence of unguided non-invasive SD detection.

Recently a case study has been reported of spreading depression correlates in the EEG delta band (0-4 Hz) visually identified in retrospectively examined 12-hr time-compressed scalp EEG records that were time-associated with depth electrode ECoG suppressions and DC shifts of a SD [115], suggesting another possible method of non-invasive SD detection.

### 4.3 SD modeling, trajectories, and simulation

Although tracking [116], modeling [117, 118], and observational [119] studies of SDs have been made in or used gyrencephalic brains, our model of an SD propagating on a flat sheet replicates the brain surface electric field of an SD appropriately and does represent the most likely situation of SD propagation along a gyrus, based on Woitzik et al.’s [41] data that 17 out of 19 SDs observed in human malignant hemispheric stroke patients travelled on gyral ridges, and on the reasoning that a gyrus can be represented as a segment of a flat sheet. One effort by Chamanzar et al. [120] provides simulations for detecting SDs based on modeling their EEG suppressions in the clinical EEG frequencies with silent cortical dipoles imbedded in a realistic brain model. This work used the well-known variation in EEG suppression duration and width as a determinant of detectability. This effort is clearly complementary to our work concerning the brain surface electric field produced from the DC-shift associated with SD depolarization, its propagation to the scalp and reconstruction as a heat-map image of the associated scalp electric field: Both SDs DC-shift, as modeled in this work, and EEG suppression, as modeled by Chamanzar et al. [120], are recognized as essential markers of SD identification using the invasive ECoG method [108].

### 4.4 Velocity and Size Errors Using Estimates from a Limited Electrode Array

The relative position of the SD focus and the electrode array affect the prediction of the size and speed of the SD. Electrodes nearest to the focal point of the SD showed the most significant errors in the velocity calculation (> 100%). This is because directly above the focus, the scalp voltage is affected by the entire ring. This effect becomes less pronounced as the ring expands outwards. The apparent velocity approaches the expansion velocity of the SD as the leading edge (scalp) propagates away from the focal point (temporally and spatially). Electrodes further from the focus had relatively small errors (∼6%) but increased (by 20%) as the electrode-sensor row was offset from the radial projection of the SD focus. Despite the large variability in velocity predictions, the thickness of the SD was more reliably calculated with errors of approximately 4 - 10% or 0.2 - 1.0 cm. The model showed that the post-SD scalp voltage recovered at a slower rate due to the same axisymmetric influence of the ring structure with the more closely spaced electric field lines at the trailing edge of the ring. This sustains the decreased voltage of the depolarization for a longer duration (t_d_) at this trailing edge. These effects may not be indicative of the truly active system, where biofluidic pathways, i.e., blood and CSF, may serve to restore voltage potentials more efficiently than a passive material. For example, blood flow can increase post-SD [41] and may remove excess ions [121; page 156]. In addition, massive amounts of K^+^ appear in the CSF [122] and their clearance via the glymphatic route [123] could alter the voltage potential.

### 4.5 Estimated Surface Characteristics and Voltage Ratio - Comparison of Simulated Square and Triangle Waves

The simulated DC-potential on the scalp has identifiable shape and magnitude characteristics. The spatial extent of the voltage change due to the SD wave expands considerably, from 2.4 mm to 30.8 mm, as it passes through tissue, allowing it to be detected by multiple electrodes simultaneously. The voltage data presented here are discussed in terms of V_ratio_, the peak scalp electrode sampled voltage scaled by the peak voltage of the SD. The simulation predicted voltage ratios of 0.18 and 0.0735 for the square wave and triangular wave, respectively. Aesthetically, the triangle wave offers a better fit to the waveforms observed by Drenckhahn et al. [34], but would result in less voltage at the scalp than the concussive-SD observed by Pacheco et al. [26; Figure 4H top], suggesting that it provides a conservative estimate of SD-detectability. The primary reason that the square wave shows better propagation of voltage (higher |V|) is that it produces more power (V^2^/R) for a given resistance.

### 4.6 Comparison of Voltage Ratios – Simulation vs. Clinical Research

The median value of the voltage ratio from the retrospective analysis of data from Drenckhahn et al. [34] was 0.0316 (0.0221, 0.0527), which is significantly different from our voltage ratio value of 0.0735 (p < 0.001). This difference is within the same general range and is acceptable for our purposes, especially because data from human concussive SDs is not available. The higher voltage ratio from our simulation of an expanding ring concussive SD can be attributed to the different shape, size, and propagating configuration of its electric field as compared to that of an aSAH SD. The strength of signal predicted at the scalp surface is stronger for an expanding ring SD compared to a globular SD because the contributions of the electric field from adjacent portions of the expanding ring add together to produce a net increase in the electric filed at the scalp. Analytic models of the expanding ring SD and an aSAH SD-like globular-region model were compared. The globular-SD had a triangular voltage profile with a width of 3 mm. These analytic models allow predictions of the scalp surface DC-voltage ratios, which are 0.093 for the expanding ring, compared to 0.010 for the globular-region model with triangular waveform. This analysis explains why our simulated concussive SD voltage ratio of 0.0735 is much larger than the ratio from our retrospective data analysis of human aSAH SDs. Furthermore, there are inherent limitations in calculating a ratio of human voltage measurements due to misalignment of the subdural electrode strip and the scalp electrodes. In the human case, there are also complications in the estimation of an SD’s maximum voltage as sensed by a limited one-dimensional ECoG electrode array and the widely separated scalp EEG electrodes in relation to an SD’s trajectory, especially as the effect of fresh craniotomy in these patients with aneurismal SAH is unknown [34].

### 4.7 SD Detectability and Practicality

Because the conductivity equation is linear and material properties are constant, the relative voltage is independent of the peak voltage (ΔV_SD_), but not the voltage waveform used to model the SD. Therefore, a single numerical experiment can be used to evaluate a full range of voltages. If we apply our simulated triangle wave voltage ratio of 0.0735 to the minimum voltage of the retrospectively determined brain-surface voltages from Drenckhahn et al. [34] of -800 µV, we obtain an estimated scalp voltage of -59 µV. This voltage is within the sensitivity threshold of currently available DC-coupled biosignal amplifiers. This suggests that the threshold of sensitivity for DC-EEG is an important consideration and that all but the weakest scalp voltages from a SD would be detectable with the caveat that noise and artifacts might interfere. It also might be possible that an SD could be non-invasively detected with an algorithm without visualization with greater than 1 cm electrode spacing.

The practical implementation of non-invasive SD detection for concussion will depend on the development of a non-invasive method in a known source of SD’s in severe acute brain injury patients in the Neurointensive Care Unit. The initial phase of this development cycle has been reported using an 8×6 cm 1-cm spaced 29 electrode array [124, 125] placed over the edge of the lesion.

### 4.8 Limitations

The SD spreads uniformly in the simulation since there is no biochemical, physical, or anatomical variation in the simulated (lissencephalic) cortex [119, 126] or in the modeled material layers that would serve to distort the shape of the wave, thus our simulation results have limited applicability to those SDs that might enter sulci of a gyrencephalic cortex and to situations where there are anatomical variations in the layers. However, the simulation suggests the design parameters of a preliminary sensor array can be used to obtain the experimental results necessary to refine the numerical model and SD detection algorithms that could be developed for such a scalp SD sensor unit. Another limitation of this current study is that the effect oof local drift such as the TEP [127, 128] and artifacts resulting from improper electrode-skin contact were not included. Another limitation applies to using our results to predict SD detection success of different SD widths and morphologies. If a globular SD, such as is typical for ischemic stroke, were modeled travelling along the edge of a lesion or separated from the lesion edge [129], a low error image reconstruction and faithful visualization might require different electrode spacing.

A sensor array with electrode spacings of 1.0 cm will enable, and is sufficient for, the reconstruction of an image of the concentrically spreading electric field on the scalp from a brain-surface SD with low error and high visual fidelity. Obviously, the addition of large-scale background noise and/or artifacts will hinder the reconstruction of the signal, as will the local failure of a single electrode depending on the overall effectiveness of the interpolation algorithms used to process the incoming signals. Other issues to be considered include the relative electrode sensor sensitivities and relative motion to the scalp.

Non-invasive SD detection is often stated as a goal in those publications where SDs are detected in acute brain injury patients using invasive ECoG [34-36], but applying our simulation results for the detection of concussion-SDs, with this being in far future, directly to acute brain injury SDs might be inappropriate, as suggested by our comparison of voltage ratios between our simulated expanding ring SDs to the SAH-SD data gleaned from Drenckhahn et al.’s work [34]. This is because our results are based on the expanding ring SD morphology, whereas SDs in acute brain injury are most likely globular with variable trajectories and dimensions including width as suggested by the SD trajectory data of Woitzik et al. [41] in malignant hemispheric stroke patients.

### 4.9 Conclusion

This simulation study suggests that propagating SDs can be visually detected on the scalp with properly configured DC-EEG. If our simulation results were used to implement successful non-invasive SD scalp detection, the link between SD and concussion could be confirmed. The confirmation of this link would involve a large clinical trial using a deployable non-invasive SD detection system that could be placed on an athlete’s head at or near the concussive focus starting within 10 minutes of the suspected concussion for a data collection period of at least 20 minutes. SD could then provide a unifying concept for the wide-ranging symptoms of concussion; the varying symptoms used to diagnose concussion could be a result of SD occurring in different cortical locations, depending on where the head-hit occurred. In addition, our results imply that the noninvasive detection of SDs in severe acute brain injury patients in the Neurointensive Care Unit is possible with the proper electrode array configuration. This implication is supported by our proof-of-concept validation data [124, 125].

## Acknowledgements

We thank Thomas Ferguson, Richard Kraig, Douglas Smith, and Joel Greenberg for reading and commenting on the manuscript and Hiba Al-Ashtal for editorial services.

## Compliance with Ethical Standards

All ethical standards have been met. See section “Retrospective Evaluation of Human Data”.

## Intellectual Property Disclosures

Parts of this work are disclosed in U.S. Patent No. 10,028,694, published and publically available on May 26, 2016 and issued July 24, 2018; and in U.S. Published Patent Application No. 2019/0038167, published February 7, 2019.

## Figure Captions

**Movie S1 (see mpg file).**
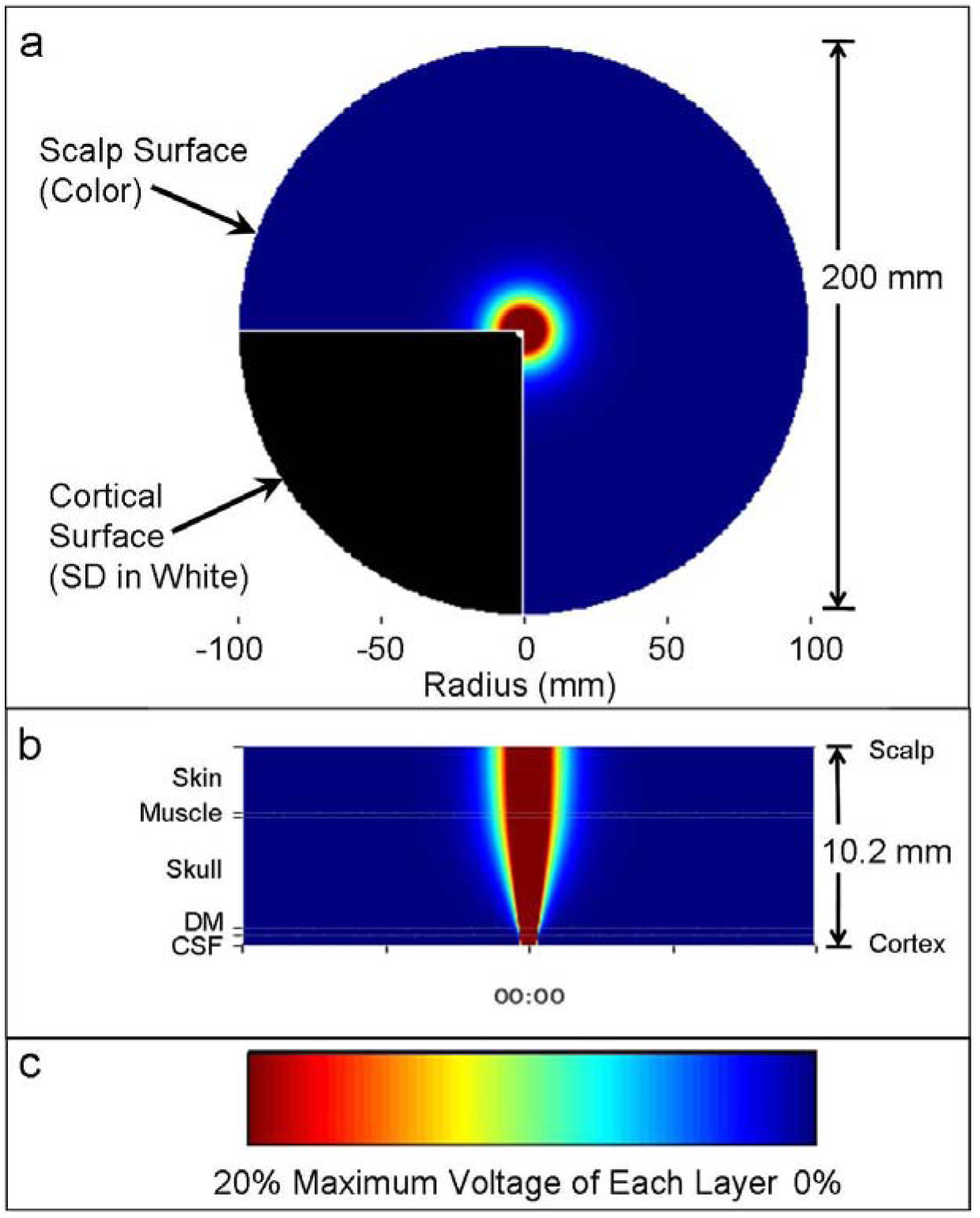
Movie of a simulation of an expanding ring cortical Spreading Depolarization (SD) showing: (a) a top view of the simulated propagating scalp voltage, represented by a 200-mm diameter disk and mapped to the pseudo-color scale, between 5 seconds and 15 minutes from the concentrically expanding SD. The 3-mm wide, expanding ring SD on the cortical surface is shown in white against black in the lower left quadrant. (b) A radial section, or cross-section of a side view, showing the propagating SD through the 5 layers [skin, muscle, skull, dura mater (DM), and cerebrospinal fluid (CSF)] from the brain surface to the scalp (thickness 10.2 mm) with a timer below. (c) The pseudo-color scale showing with the voltage scaled to 20% of the maximum voltage of each layer

**Supplementa1 Fig. 1.**
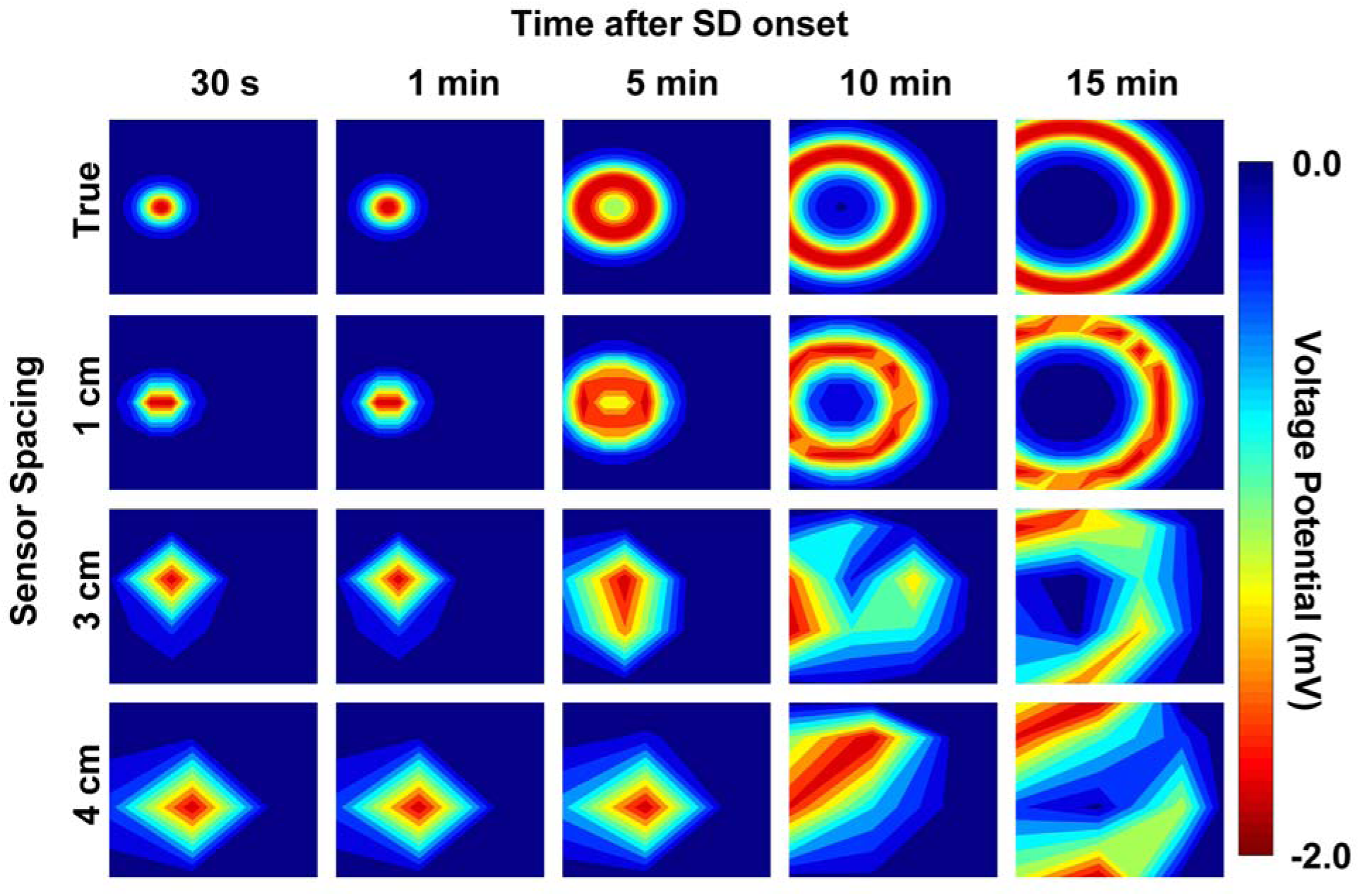
The scalp voltage intensity maps from a cortical spreading depolarization (SD) onset as a function of sensor spacing and time after onset. Top row: The exact voltage field is provided for reference at 0.5, 1, 5, 10, and 15 minutes after SD initiation. The bottom three rows show the simulated voltages at the scalp for the electrode spacings of 1, 3, and 4 cm

